# Susceptibility of midge and mosquito vectors to SARS-CoV-2 by natural route of infection

**DOI:** 10.1101/2020.09.29.317289

**Authors:** Velmurugan Balaraman, Barbara S. Drolet, Natasha N Gaudreault, William C. Wilson, Jeana Owens, Dashzeveg Bold, Dustin A. Swanson, Dane C. Jasperson, Leela E. Noronha, Juergen A. Richt, Dana N Mitzel

## Abstract

SARS-CoV-2 is a recently emerged, highly contagious virus and the cause of the current pandemic. It is a zoonotic virus, although its animal origin is not clear yet. Person-to-person transmission occurs by inhalation of infected droplets and aerosols, or by direct contact with contaminated fomites. Arthropods transmit numerous viral, parasitic, and bacterial diseases; however, the potential role of arthropods in SARS-CoV-2 transmission is not fully understood. Thus far, a few studies have demonstrated that SARS-CoV-2 replication is not supported in cells from certain insect species nor in certain species of mosquitoes after intrathoracic inoculation. In this study, we expanded the work of SARS-CoV-2 susceptibility to biting insects after ingesting a SARS-CoV-2infected blood meal. Species tested included *Culicoides sonorensis* biting midges, as well as *Culex tarsalis* and *Culex quinquefasciatus* mosquitoes, all known biological vectors for numerous RNA viruses. Arthropods were allowed to feed on SARS-CoV-2 spiked blood and at various time points post infection analyzed for the presence of viral RNA and infectious virus. Additionally, cell lines derived from *C. sonorensis* (W8a), *Ae. aegypti* (C6/36), *Cx. quinquefasciatus* (HSU), and *Cx. tarsalis* (CxTrR2) were tested for SARS-CoV-2 susceptibility. Our results indicate that none of the biting insects, nor the insect cell lines support SARS-CoV-2 replication. We conclude, that biting insect do not pose a risk for transmission of SARS-CoV-2 to humans or animals following a SARS-CoV-2 infected blood meal.

## Introduction

Severe acute respiratory syndrome coronavirus 2 (SARS-CoV-2) is the causative agent of the 2019 coronavirus disease (COVID-19) pandemic. SARS-CoV-2 belongs to the order *Nidovirales*, family *coronaviridae*, and genus *betacoronavirus*. It is an enveloped virus with a positive-sense, single-stranded RNA genome of approximately 30 kb in length (Chen et al. 2020). SARS-CoV-2 infects humans, and has the potential to infect various animal species (Chu et al. 2020). Transmission from these animals to humans is not yet clearly understood. The virus is mainly transmitted from person-to-person by inhalation of droplets and aerosols produced by infected people (Chan et al. 2020), or through contact with contaminated surfaces (Goldman 2020, Sonja A. Rasmussen 2020; Kwon et al., 2020). Arthropods transmit numerous diseases to humans and animals via biological and mechanical transmission (Leitner et al. 2015). Although the SARS-CoV-2-related coronaviruses SARS-CoV-1 and MERS-CoV are not transmitted by insects, concerns have been raised, by those in both public health and agricultural sectors, as to their potential role in spreading SARS-CoV-2 among humans and animals. For arthropods to be transmission-competent vectors, the respective pathogen must be acquired from a host during blood feeding, then infect the midgut, escape the midgut barrier, disseminate to and infect the salivary glands, and finally be transmitted to a susceptible host during subsequent blood feeding (Franz et al. 2015). A recent report demonstrated that SARS-CoV-2 replication was not supported in *Aedes aegypti, Ae. albopictus* and *Culex quinquefasciatus* mosquito species after an intrathoracic route of infection (Huang et al. 2020). Another report showed that the SARS-CoV-2 does not replicate in cells derived from *Aedes* mosquitoes, nor was it present in field-caught *Culex* and *Anopheles* mosquitoes from Wuhan (Xia et al. 2020).

Here, we report the first susceptibility study of SARS-CoV-2 infection using three critical insect vectors following ingestion of a SARS-CoV-2 infected blood meal, including an agriculturally important animal disease vector, *Culicoides sonorensis* biting midges, and two significant human disease vector mosquito species, *Cx. tarsalis* and *Cx. quinquefasciatus*. Additionally, four insect-derived cell lines from *C. sonorensis* (W8a), *Ae. aegypti* (C6/36), *Cx. tarsalis* (CxTrR2), and *Cx. quinquefasciatus* (HSU) were also evaluated for SARS-CoV-2 susceptibility.

## Methods

The SARS-CoV-2 USA-WA1/2020 strain was acquired from Biodefense and Emerging Infection Research Resources Repository (BEI Resources, Manassas, VA, USA) and was passaged three times on VeroE6 cells (ATCC, VA, USA) with a final titer of 2.5 x 10^6^ TCID_50_/ml. Arthropod cell cultures were derived from *C. sonorensis* embryos (W8a; McHolland and Mecham 2003), *Cx. tarsalis* embryos (CxTrR2; Arthropod-Borne Animal Diseases Unit; ABADRU, Manhattan, KS, USA), *Cx. quinquefasciatus* ovaries (HSU; Hsu et al. 1970), and *Ae. albopictus* larva (C6/36). The W8a, CxTrR2, HSU, and C6/36 cells were maintained in CuVa medium, L-15 medium (with 10% tryptose phosphate broth) and Medium 199H, respectively. All media (Sigma-Aldrich, St. Louis, MO, USA) was supplemented with 10-20% FBS (ITFBS; Sigma). Cells were maintained at 27°C in closed T-flasks and inoculated with SARS-CoV-2 at approximately 0.1 multiplicity of infection (MOI) for 1h before the inoculum was replaced with fresh culture media. Cell cultures were monitored for cytopathic effect (CPE) by light microscopy and culture supernatants were collected at 0, 2, 4, and 8 days post infection (dpi) for subsequent titration by TCID_50_-CPE assay on VeroE6 cells.

*Cx. tarsalis, Cx. quinquefasciatus*, and the ABADRU *C. sonorensis* colonies were reared and maintained in the ABADRU insectary. Arthropods were transported to Kansas State University, Biosecurity Research Institute (BRI) for infection studies under Arthropod Containment Level-3 (ACL-3) conditions.

Adult female *C. sonorensis* (n=200) midges were allowed to feed on defibrinated sheep blood mixed 1:1 (v/v) with SARS-CoV-2 (2.0 x 10^6^ TCID_50_/ml). Negative control unfed midges (n=100) were maintained in adjacent cages. For mosquitoes, 8-day old *Cx. tarsalis* (n=100) or 10-day old *Cx. quinquefasciatus* (n=100) were allowed to feed on SARS-CoV-2 spiked sheep blood as described above. Negative control mock-infected blood-fed *Cx. tarsalis* (n=50) were maintained in adjacent cages. After an hour of feeding, midges or fully engorged mosquitoes were held at 28°C for 10 days. Surviving midges and mosquitoes at day 10 were pooled (n=5-10) in 1 ml virus transport media (199E media supplemented with antibiotic-antimycotics; Sigma), and stored at −80°C until processed for virus isolation (VI) and RNA extractions.

Pooled arthropods were homogenized by a Tissuelyser II (Qiagen, Germantown, MD, USA) using tungsten carbide beads (Qiagen). An aliquot (140 μl) of homogenate was used for RNA extraction with the remaining homogenate filtered through a 0.22 μm PES membrane filter (MIDSCI, St. Louis, MO, USA) before subsequent VI.

RNA extraction was performed using the QIAamp viral RNA mini kit (Qiagen) as per manufacturer’s instructions. RT-qPCR assay was performed according to the Center for Disease Control (CDC) protocol for detection of SARS-CoV-2 nucleocapsid (N)-specific RNA (https://www.fda.gov/media/134922/download) using Script XLT One-Step RT-qPCR Tough Mix (Quanta Biosciences, Beverly, MA, USA) on a CFX96 Real-time thermocycler (BioRad, Hercules, CA, USA). Plate controls included a quantitated SARS-CoV-2 N-specific qPCR positive control, diluted 1:10 (Integrated DNA Technologies, IA, USA), and a non-template control (NTC). Results were analyzed using the Bio-Rad CFX Manager 3.1 with samples below 40 Ct considered positive.

Arthropod homogenates (100 μl) were added on to VeroE6 cells in 24 well plates and incubated at 37°C and 5% CO_2_ for 3 days. Culture supernatents were blind-passaged three times on VeroE6 cells, and at the first and third passage, cells were examined by an indirect immunofluorescence assay (IFA) for the presence of SARS-CoV-2 antigen. Briefly, 3 dpi cells were fixed with ice cold 100% methanol for 10 mins at −80°C and washed three times with 1 x PBS Tween 20 (0.05%). Mouse monoclonal antibodies (in house) specific for the Receptor Binding Domain (RBD) of spike protein of SARS-CoV-2 was diluted 1:5 in 1x PBS containing 1% BSA and 150 μl was added to each well and incubated at room temperature (RT) for 1 h. The cells were washed three times as described above, and then incubated with 150 μl of FITC-conjugated goat anti-mouse IgG (Abcam, Cambridge, MA, USA), diluted 1:500 in 1x PBS with BSA, for 1 h at RT. After washing and drying, cell monolayers were examined by an EVOS fluorescent microscope (ThermoFisher Scientific, Waltham, MA, USA) for the presence of FITC positive cells. Mock infected and SARS-CoV-2 infected VeroE6 cells were used as negative and positive controls, respectively.

## Results and discussion

The goal of this study was to determine whether arthropods are susceptible to SARS-CoV-2 by a natural route of infection which has not yet been evaluated. In addition to important mosquito vectors, *Culicoides* midges were also evaluated in this study. Initial infection studies were performed *in vitro* with the insect cell lines W8a, C6/36, CxTrR2, and HSU. Two independent experiments showed no obvious sign of CPE for any of the SARS-CoV-2-infected arthropod-derived cell cultures, nor for any of the insect culture supernatants collected at 2, 4 or 8 dpi and passaged on VeroE6 cells.

Next, susceptibility of insects after an infectious blood meal was investigated. Of 200 midges allowed to feed on the SARS-CoV-2-spiked blood meal, 140 survived until 10 days post blood meal and were further analyzed. The majority (85%) of virus-fed midge pools had detectable SARS-CoV-2-specific RNA with an average Ct value of 34.84±2.6; the day 10 control unfed midges were negative for SARS-CoV-2 RNA (Table 1). Among the *Cx. tarsalis* mosquitoes allowed to feed on SARS-CoV-2-spiked blood (n=100), only 48 virus-fed mosquitoes survived until day 10. One out of 6 (17%) virus-fed *Cx. tarsalis* mosquitoes had detectable SARS-CoV-2-specific RNA with an average Ct value of 31.3, and none of the 30 mock-infected blood-fed *Cx. tarsalis* mosquitoes were SARS-CoV-2 RNA positive (Table 1). Similarly, of 100 *Cx. quinquefasciatus* mosquitoes, 47 virus-fed survived until day 10. Viral RNA was detected in 50% of the SARS-CoV-2-fed mosquitoes with an average Ct value of 34.17 (Table 1).

**Table 1:** Detection of SARS-CoV-2 viral RNA in various arthropods by RT-qPCR

| Treatment Group | Positive homogenate pools out of total (% positive) | Mean Ct ± SD |
| --- | --- | --- |
| Unfed *C. sonorensis* | 0/4 (0) | ND* |
| SARS-CoV-2 fed *C. sonorensis* | 12/14 (85) | 34.84±2.6 |
| Mock-infected fed *Cx. tarsalis* | 0/3 (0) | ND* |
| SARS-CoV-2 fed *Cx. tarsalis* | 1/6 (17) | 31.3 |
| SARS-CoV-2 fed *Cx. quinquefasciatus* | 3/6 (50) | 34.17±3.3 |

\*ND=not detected

To determine the presence of infectious virus, serial passages of pooled arthropod homogenates were performed on VeroE6 cells. No CPE was observed after three passages of virus-fed *Culicoides* midge homogenates, and IFA analysis of passage one and three of inoculated VeroE6 cells confirmed the absence of SARS-CoV-2 (Table 2; Figure 1). SARS-CoV-2-infected VeroE6 cells were used as an IFA positive control and showed a clear positive staining pattern (Fig.1). Unfed control midge samples were negative by VI and IFA (Table 2). Similarly, no infectious virus was detected from any of the six homogenate pools of SARS-CoV-2-fed *Cx. tarsalis* mosquitoes that were passaged on VeroE6 cells and analyzed for CPE and by IFA; the control *Cx. tarsalis* homogenate pools were also negative by both methods (Table 2). The six virus-fed *Cx. quinquefasciatus* mosquito homogenates tested for infectivity by VI and IFA were also negative for SARS-CoV-2 (Table 2).

**Figure 1.**
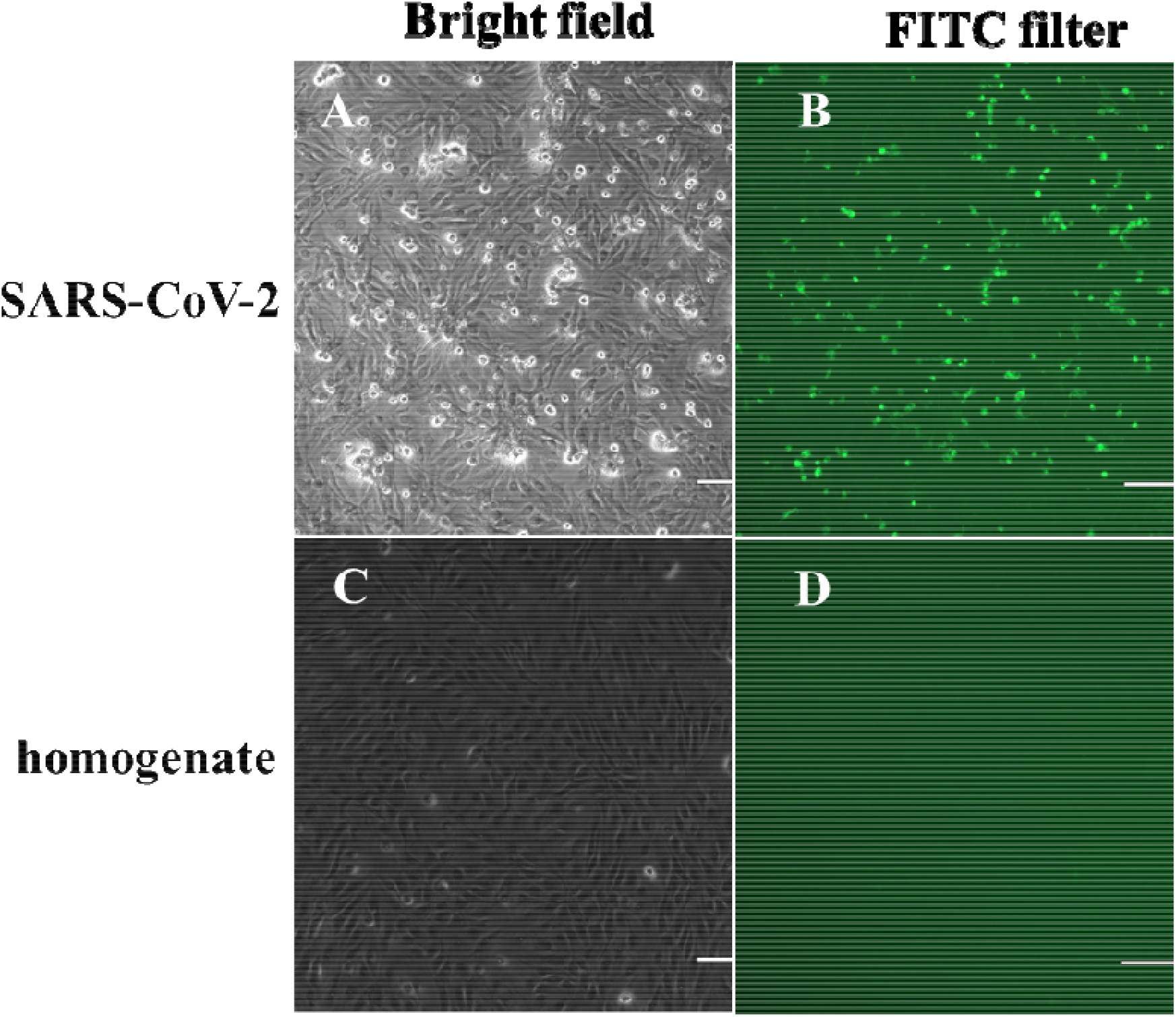
Indirect immunofluorescence assay for the detection of SARS-CoV-2 infected cells. SARS-CoV-2 and a representative passage 3 arthropod homogenate were incubated 3 days on VeroE6 cells. A) Bright field; B) Positive SARS-CoV-2 infected VeroE6 cells (positive control cells); and C) Bright field; D) VeroE6 cells inoculated with passage three of a RT-qPCR+ *Culicoides sonorensis* day 10 homogenate. (Magnification 10x)

**Table 2:** Detection of infectious SARS-CoV-2 in arthropod homogenate pools on VeroE6 cells by virus isolation and an indirect immunofluorescence assay

| Species | Control arthropods |  | SARS-CoV-2 fed arthropods |  |
| --- | --- | --- | --- | --- |
|  | VI<br>(positive/N) | IFA<br>(positive/N) | VI<br>(positive/N) | IFA<br>(positive/N) |
| *C. sonorensis* | 0/4 | 0/4 | 0/14 | 0/14 |
| *Cx. tarsalis* | 0/3 | 0/3 | 0/6 | 0/6 |
| *Cx. quinquefasciatus* | N/A | N/A | 0/6 | 0/6 |

N= total number of day 10 homogenate pools. 0= no virus positive or FITC positive cell cultures; N/A = not tested

A key factor crucial for arthropod-mediated transmission is that the infected host, either person or animal, is viremic at the time of feeding. Thus far, SARS-CoV-2 is known to cause viremia in some cases of infected people (Young et al. 2020). Susceptible animal models tested so far appear to be aviremic (Shi et al. 2020) except for hamsters which regularly show viremia (Chan et al. 2019). Therefore, controlled studies to rule out arthropod transmission of this RNA virus are critical for determining risk, as well as developing accurate epidemiological modeling and control strategies.

Overall, our results agree with previously published findings that *Aedes* mosquito derived cells do not support SARS-CoV-2 replication (Xia et al. 2020). Additionally, we have shown that two different *Culex* species derived cell lines and one *Culicoides* midge derived cell line are also refractory to SARS-CoV-2 infection. In a previously published SARS-CoV-2 susceptibility study in mosquitoes, intrathoracic injection of SARS-CoV-2 grown in Vero76 was used to determine the susceptibility of *Ae. aegypti, Ae. Albopictus*, and *Cx. quinquefasciatus* to the virus (Huang et al. 2020); however, intrathoracic inoculation bypasses the natural route of infection via the ingestion of a virus-infected blood meal (Franz et al. 2015). Therefore, in the present study, testing of arthropods for SARS-CoV-2 susceptibility was performed following the ingestion of a virus-spiked blood meal; this is important not only because it is the most relevant route of infection for arthropod vectors, but also because of the quasi-species nature of an RNA virus inoculum such as for SARS-CoV-2 (Jary et al. 2020). Considerable genetic bottle necks and natural selection processes exist for viruses in arthropod replication and arthropod-borne transmission. Depending on the insect vector, the number of virus particles ingested via a blood meal depends on the level of viremia in the host and the volume of the meal. When virus particles ingested via the blood meal enter the midgut, few will be able to infect the midgut epithelium. After midgut replication, progeny viruses escape the midgut barrier and disseminate in the insect’s hemocoel, and then next they infect and replicate in the salivary gland. This is critical for subsequent virus transmission to a susceptible host during a blood meal. The quasis-pecies nature of RNA viruses combined with the natural selection for defined viral genotypes enables RNA viruses to infect and replicate in arthropods and results in a defined virus population selected for increased fitness for the arthropod environment. These viral genotypic changes could lead to viral biotypes with a higher ability of salivary gland infection and bite transmission, when compared to a virus which is artificially injected into the insect’s hemocoel.

Our *in vitro* and *in vivo* studies of midge and mosquito susceptibility to SARS-CoV-2 infection following a natural ingestion route of exposure showed that viral RNA remained in virus-fed arthropods for up to 10 days post virus-spiked blood feeding. However, no infectious virus was recovered from these RNA-positive arthropods, even after three passages on highly susceptible VeroE6 cells. Our in vitro studies using various insect cells support these results since no infectious virus was detected in supernatants of SARS-CoV-2 inoculated insect cell cultures. In conclusion, the insect vector species known to transmit animal and human pathogens used in this study are refractory to SARS-CoV-2 infection under experimental conditions and, therefore, most likely do not play a role in transmission of SARS-CoV-2.

## Acknowledgments

We gratefully thank the staff of KSU Biosecurity Research Institute. The following reagent was obtained through BEI Resources, National Institute of Allergy and Infectious Diseases (NIAID), National Institutes of Health (NIH): SARS-CoV-2 Virus strain USA-WA1/2020 (catalogue # NR-52281). C6/36 cells were kindly provided by Robert B Tesh, UTMB, Galveston, TX.*Cx. tarsalis* and *Cx. quinquefasciatus* colonies were kindly provided by Chris Barker and O. Winokur, University of California-Davis.

## Conflict of interest

All the authors declare no conflict of interest.

## Disclaimer

The conclusions in this report are those of the author and do not necessarily represent the views of the United States Department of Agriculture.

## Funding

Funding for this study was in part by the United States Department of Agriculture (BSD, DM, DS, JO, DCJ, LN, and WCW) and through grants from NBAF Transition Funds and KSU internal funds to JAR. This study was also partially supported by NIAID Centers of Excellence for Influenza Research and Surveillance (CEIRS; contract #HHSN 272201400006C), and the Department of Homeland Security Center of Excellence for Emerging and Zoonotic Animal Diseases (grant #2010-ST061-AG0001) to JAR.

## Notes

### Competing Interest Statement

The authors have declared no competing interest.

